# Intranasal oxytocin increases early tactile engagement with a robotic dog: A within-session effect without consolidation

**DOI:** 10.64898/2026.11.20.768097

**Authors:** Seval Aydin, Kelsey C. Mitchell, Mary R. Lee, Martin O’Hara, Jonathan Gratch, Ewart J. de Visser, Hasan Ayaz, Xingnan Li, Adrian A. Franke, Frank Krueger

**Affiliations:** School of Systems Biology, George Mason University, Fairfax, Virginia, United States of America; Psychiatric Neuromodulation Service, Veterans Affairs Medical Center, Washington, District of Columbia, United States of America; Virginia Hospital Center, Arlington, Virginia, United States of America; Institute for Creative Technologies, University of Southern California, Los Angeles, California, United States of America; Warfighter Effectiveness Research Center, United States Air Force Academy, Colorado Springs, Colorado, United States of America; School of Biomedical Engineering, Science and Health Systems, Drexel University, Philadelphia, Pennsylvania, United States of America; University of Hawaii Cancer Center, Honolulu, Hawaii, United States of America

## Abstract

Oxytocin (OXT) is centrally involved in mammalian affiliation, but whether it modulates affiliative behavior toward artificial social agents, and how such an effect unfolds within a single interaction, is unclear. In a randomized, double-blind, placebo (PLA)-controlled study, 61 healthy adult males received 40 IU intranasal OXT or PLA before interacting with a Sony aibo robotic dog across an initial free interaction, a cooperative task-learning phase, and a final free interaction, with ten affiliative behaviors coded from video. OXT increased tactile engagement early: participants touched aibo longer during the initial free interaction, the only outcome of ten to survive family-wise correction. A weaker increase in smile frequency did not survive correction, and instrumental behavior was unaffected. Contrary to our second prediction, the advantage was not consolidated: it held through the initial interaction, closed across the intervening task, and was absent at the final one. An exploratory decomposition located the effect: participants under OXT did not initiate contact more often, and the difference was carried by how long each episode of contact was sustained. Urinary OXT confirmed group-level exposure. Because the design included no comparison condition and confounded interaction period with elapsed time and pharmacokinetics, it cannot establish bond formation, robot specificity, or consolidation as a psychological process. What it does bound is where and when OXT acted: on the maintenance of contact rather than its initiation, and only while the agent was novel—a pattern that fits salience- and novelty-based accounts better than bond formation.

## 1. Introduction

The neuropeptide oxytocin (OXT) is centrally involved in mammalian affiliation, yet its effects on human social behavior are neither uniformly prosocial nor context-independent. The social salience hypothesis proposes that OXT modulates the salience of socially meaningful cues in a manner contingent on context, individual differences, and interaction goals [1,2]. An allostatic perspective further suggests that OXT’s effects may be strongest under conditions of novelty and uncertainty, diminishing as environments become more predictable [3]. A central implication of these accounts is that OXT’s effects on affiliation should be strongest early in an encounter, while a partner is novel and its social status unresolved, and should weaken as the interaction becomes familiar or predictable. A key unresolved question for social neuroendocrinology is whether these OXT-modulated affiliative processes extend to artificial agents that provide social cues but lack the biological reciprocity of living social partners. The present study addressed this question by testing whether intranasal OXT modulates affiliative behavior toward a dog-like robotic agent, and whether any such effects are temporally bounded by interaction context [4–8].

Robot-dog embodiments such as the Sony <u>A</u>rtificial Intelligence Ro<u>bo</u>t (aibo) provide a useful boundary condition. Aibo offers animal-like social cues that may invite affiliative interpretation, yet lacks the reciprocal biological signaling that normally sustains affiliation with living partners [9,10]. Dog–human bonding is among the best-characterized interspecies affiliative systems, with OXT-linked positive feedback loops involving gaze and affiliation [11]; however, urinary OXT increases across repeated interactions with real dogs but progressively decreases with aibo [12], suggesting that sustained social-partner attribution may be critical for maintaining affiliative neuroendocrine dynamics. Aibo is therefore treated here not as equivalent to a dog but as a socially ambiguous artificial partner: dog-like enough to invite affiliation, artificial enough to test whether OXT-modulated engagement can emerge without biological reciprocity.

Direct tests of whether intranasal OXT shapes affiliative behavior toward artificial agents remain sparse, although endogenous OXT has been reported to rise in people who form affiliative relationships with companion robots [13], and available evidence suggests plausibility. Intranasal OXT increases gaze toward socially diagnostic stimuli, modulates anthropomorphic interpretation of non-human stimuli, enhances socially reinforced learning and emotional empathy [14], and modulates trust in interactions with automated partners in ways that depend on anthropomorphism [6,15–18]. OXT may therefore not simply increase sociability but alter whether ambiguous cues are treated as socially relevant. Novelty-dependent modulation is supported by evidence that OXT enhances attentional engagement to novel stimuli, with effects diminishing across repeated exposures [16]. Effects toward artificial agents should thus be most likely during early encounters, when the agent is novel and socially ambiguous, and may weaken as the interaction becomes familiar or instrumental. Intranasal administration remains the dominant approach for probing OXT’s causal role in human social behavior, though uncertainty about dose-response relationships, assay reliability, statistical power, and peripheral-central correspondence warrants caution [19–25].

The present study tested whether intranasal OXT enhances early nonverbal affiliative engagement toward a socially ambiguous artificial agent, characterized the behavioral specificity of any such effect, and determined whether it persists as the interaction unfolds. We used a randomized, double-blind, PLA-controlled design in which participants interacted with aibo across three sequential phases, and quantified affiliative engagement across duration, frequency and the temporal organization of behavior in several channels [26,27]. Hypothesis 1 (H1) predicted an early effect concentrated in tactile engagement, graded across channels and largely absent for instrumental behavior; H2 predicted that this advantage would be consolidated through the cooperative task phase, with the PLA group catching up to a maintained OXT group [28,29]. We additionally examined the fine-grained trajectory across the four 2.5-min segments, candidate processes that might account for any departure from H2, and—because total contact time cannot separate the initiation of contact from its maintenance—a decomposition of tactile engagement into the number of touch episodes and their mean duration. This decomposition was derived during analysis rather than specified in advance; it is reported as exploratory throughout, and it proved to be the most informative analysis in the study.

## 2. Materials and methods

### 2.1. Participants

A total of 65 healthy male adults aged 18–40 years were enrolled at George Mason University (GMU) and randomly assigned to receive either intranasal OXT (*n* = 32) or PLA (*n* = 33). Primary behavioral analyses were conducted in 61 participants with usable behavioral data (OXT *n* = 30, PLA *n* = 31). This sample is independent of the one reported in our earlier observational aibo study [12]; no participant took part in both. Demographic characteristics and baseline group comparisons for the analyzed sample are reported in **Table 1** and **S1 Appendix, Table A**. Participants were recruited using Institutional Review Board (IRB)-approved flyers posted on and off campus (S1 Appendix, Supplementary Methods). Participants were enrolled between April 22, 2022, and September 30, 2022.

**Table 1.** Baseline characteristics of the analyzed sample.

| **Characteristic** | **OXT (*n* = 30)** | **PLA (*n* = 31)** | **N: OXT/PLA** | **SMD** |
|---|---|---|---|---|
| Sample size | 30 | 31 | — | — |
| Age (years), *M* (*SD*) | 21.40 (4.49) | 23.90 (5.34) | 30/31 | -0.51 |
| Race, n (%) |  |  | 30/31 | — |
| White | 10 (33.3%) | 13 (41.9%) |  |  |
| African American | 1 (3.3%) | 3 (9.7%) |  |  |
| Asian | 17 (56.7%) | 11 (35.5%) |  |  |
| Hispanic | 2 (6.7%) | 4 (12.9%) |  |  |
| Believed received OXT, n (%) | 11 (36.7%) | 8 (25.8%) | 30/31 | — |
| Dog liking (1–3), *M* (*SD*) | 1.90 (0.31) | 1.90 (0.40) | 29/31 | -0.02 |
| Previous dog owner, n (%) | 18 (69.2%) | 15 (51.7%) | 26/29 | +0.36 |
| Current dog owner, n (%) | 9 (34.6%) | 9 (31.0%) | 26/29 | +0.08 |

**Note.** Values are *M* (*SD*) for continuous variables and n (%) for categorical variables. Variables with \|SMD\| > 0.25 were carried into the covariate-adjusted sensitivity analyses (S1 Appendix, Table K); the full baseline table is S1 Appendix, Table A. “Believed received OXT” is the blinding check reported in 3.5. M, mean; OXT, oxytocin; PLA, placebo; SD, standard deviation; SMD, standardized mean difference (OXT − PLA, pooled SD).

Inclusion criteria were male sex, age 18–40 years, good general health, and the ability to provide written informed consent (S1 Appendix, **Supplementary Methods**). Medical screening was conducted under the supervision of study physicians (M.R.L. & M.O.) and included medical history, a basic physical examination, blood-pressure and heart-rate assessment, and SCL-90-R evaluation (2.2). Participants proceeded with nasal-spray administration only after study physicians confirmed that medical eligibility criteria were met. All participants provided written informed consent. The study was reviewed and approved by Advarra IRB (Columbia, MD, USA; protocol Pro00034329). Before oversight transferred to Advarra, the study was approved by the George Mason University IRB. The study was conducted under Department of Defense protocol 11774A and in accordance with the Declaration of Helsinki. The study was not registered in a public clinical-trial registry. The study was a single-session behavioral pharmacology experiment with no health-related outcome, conducted under IND 110,562.

Analyses were conducted using available cases for each outcome. Participant enrollment, missing-data patterns, and analysis-specific sample sizes are reported in **S1 Appendix, Table B** and **S1 Appendix, Fig A**.

### 2.2. Stimuli and measures

#### Aibo

Six Sony aibo robotic dogs (model ERS-1000, 2018 version) were used (S1 Appendix, **Supplementary Methods**). Five units were assigned to participants during the interaction phase, one per testing stall, and a sixth unit was reserved for experimenter-led task demonstrations. For three participants the unit code in the data file does not correspond to one of the five stall units; these cases are examined in S1 Appendix, **Supplementary Results.**

#### The aibo platform

The ERS-1000 is an autonomous companion robot with touch sensors on the head, chin and back, touch-sensitive paw pads, a fish-eye camera at the nose used for face and object recognition, a second camera at the waist for spatial mapping, multiple microphones and a speaker. It responds to tactile, auditory and visual input with locomotion, posture change, head and tail movement, eye-display animations and vocalizations, so the timing of its responses was contingent on participant behavior.

#### Self-report questionnaires

Participants completed standardized measures of empathy (Interpersonal Reactivity Index, IRI) [30], attachment style (Relationship Questionnaire, RQ) [31], personality (NEO Five-Factor Inventory, NEO-FFI) [32], state and trait anxiety (State-Trait Anxiety Inventory, STAI) [33], positive and negative affect (Positive and Negative Affect Schedule, PANAS) [34], and general symptomatology (Symptom Checklist-90-Revised, SCL-90-R) [35]. Robot perceptions were assessed before and after the interaction using the Robot Likeability Measure (RLM) [36] and a study-specific 12-item robot-description scale (10DIM); attitudes toward robots and dogs were assessed with the Robot Dog Questionnaire (RDQ) [37], previously used in our laboratory [12]. Control questions assessed dog ownership history, general attitudes toward dogs, and perceived treatment assignment (2.3). All self-report instruments are summarized in **S1 Appendix, Table C**.

### 2.3. Setup and procedure

#### Experimental setting

All sessions were conducted at the Krasnow Building at GMU in a large room partitioned into five individual stalls. This setup prevented participant interaction while allowing continuous monitoring by study staff (S1 Appendix, **Supplementary Methods**). Up to five participants were tested simultaneously, and participants were assigned to stalls, and hence to aibo units, at random.

#### Experimental procedure

The study followed a randomized, double-blind, PLA-controlled, between-subjects design. Participants were randomly assigned in a 1:1 ratio to receive either 40 International Units (IU) of intranasal OXT (Syntocinon^®^, Pharmaworld, Zurich, Switzerland) or PLA according to a pre-generated block-randomization schedule prepared by F.K., who alone held the allocation key and was not involved in data collection or behavioral coding. The 40-IU dose follows prior single-dose behavioral OXT work [38–40] and established safety and dosing guidance [23,41]. Sprays were labeled by code only; the PLA consisted of the same vehicle solution without the active peptide and was matched in appearance, spray volume, and number of administrations. Participants, the experimenters who ran the sessions, the study physicians, and the behavioral coders were therefore blind to treatment assignment, and coders were additionally blind to interaction period. The allocation was not revealed until all primary and secondary analyses were complete.

The session comprised three phases: pre-interaction (∼90 min), interaction (∼30 min), and post-interaction (∼60 min); the interaction phase comprised an initial free interaction (A), a structured task phase (B), and a final free interaction (C) (**Fig 1**).

**Fig 1.**
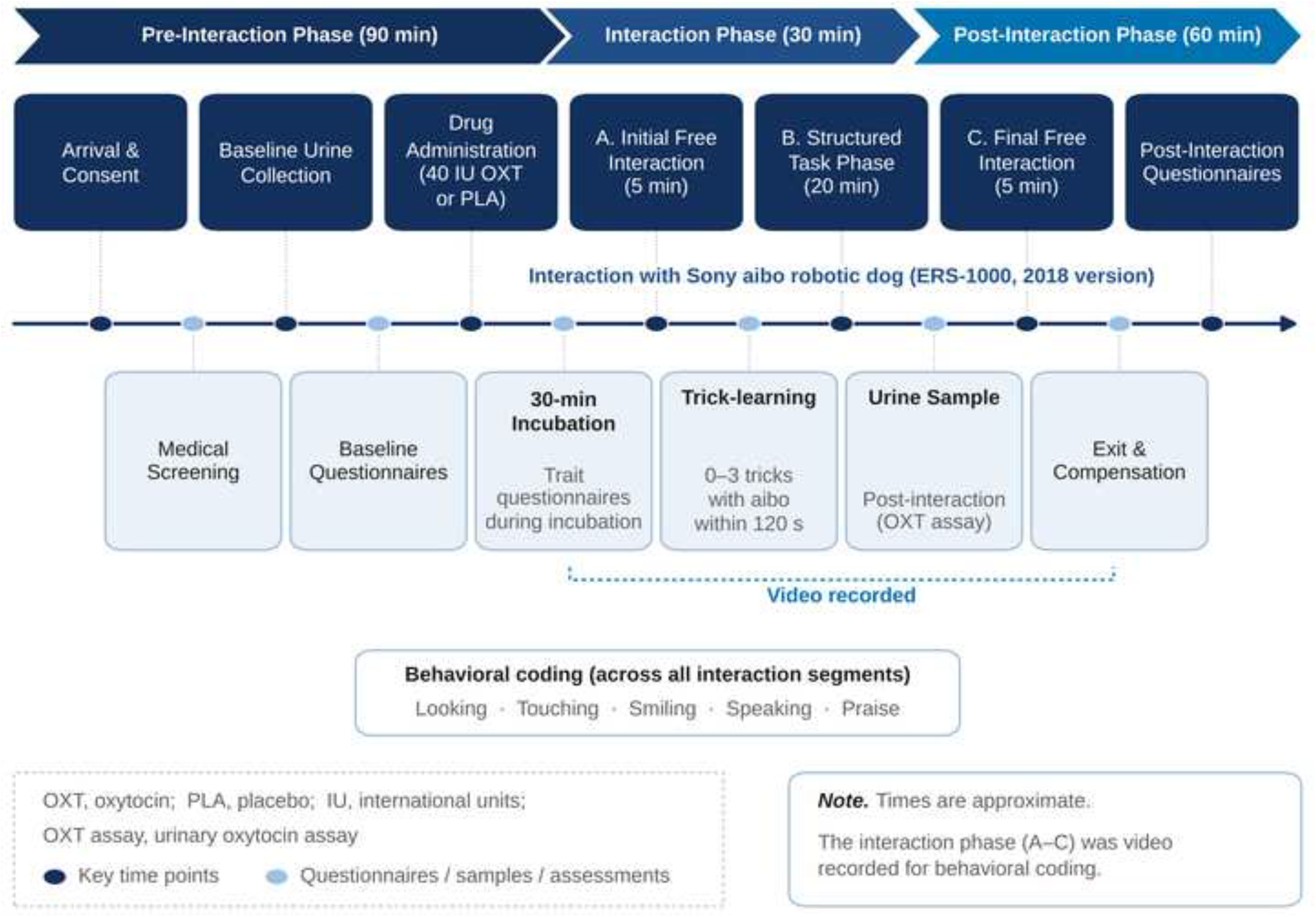
Experimental design and study timeline. Participants completed a single session comprising a pre-interaction phase (∼90 min), an interaction phase (∼30 min), and a post-interaction phase (∼60 min). After enrollment, medical screening, baseline questionnaires, and a baseline urine sample, participants received 40 IU intranasal OXT or PLA and completed a 30-min incubation period during which trait questionnaires were administered. The interaction phase comprised three consecutive segments with the Sony aibo robotic dog (ERS-1000, 2018 version). (A) Initial free interaction. Five minutes of unstructured interaction; the primary analyses are based on this segment. (B) Structured task phase. Twenty minutes in which participants attempted to teach aibo three tasks in a fixed order (squats, copy me, learn this), each within a 120-s window; only the third requires touching aibo. (C) Final free interaction. A further five minutes of unstructured interaction, matched to (A). All three segments were video recorded and later coded for looking, touching, smiling, speaking, and praise behaviors. A second urine sample was collected after the interaction phase, approximately 60 min after administration. IU, international units; OXT, oxytocin; PLA, placebo.

After medical clearance and baseline urine collection, participants self-administered the assigned spray under medical supervision (five sprays per nostril). A 30-min incubation followed—aligned with a commonly used post-administration window [23]—during which participants completed trait questionnaires (IRI, NEO-FFI, RQ) and repeated state measures (STAI, PANAS).

The video-recorded interaction phase comprised three segments: an initial free interaction (5 min), a structured task-learning phase (∼20 min), and a final free interaction (5 min).

During the initial free interaction, participants engaged freely with aibo without task constraints to capture spontaneous affiliative behavior. During the structured task-learning phase, participants completed three predefined tasks, always in the same order (S1 Appendix, **Supplementary Methods**). Each task was introduced and demonstrated by the experimenter (S.A. & K.C.M.), and participants then returned to their stalls to attempt the task with their assigned aibo for approximately 120 s. Following the structured phase, participants again interacted freely with aibo during the final free interaction, allowing assessment of changes in affiliative behavior after the interaction context had progressed through the task phase.

During the post-interaction phase, participants provided a second urine sample, completed post-interaction questionnaires (STAI, PANAS, RLM, RDQ, 10DIM), underwent a final medical check, reported perceived treatment assignment (OXT vs. PLA), and received compensation.

### 2.4. Data analysis

#### Power analysis

A sensitivity analysis indicated that the achieved sample (OXT *n* = 30, PLA *n* = 31) provided 80% power to detect a between-group effect of *d* = 0.73 (α = .05, two-tailed) at the initial free interaction. This is the smallest effect the design could reliably detect, not a prediction, and the design was not powered for the Period × Treatment interactions, the T1–T4 models, or the exploratory moderation models (S1 Appendix, **Supplementary Methods**). Reported effect sizes should therefore be read as evidence about the presence of an effect rather than as calibrated estimates of its size [42]. Non-significant results in the underpowered analyses are inconclusive.

#### Oxytocin analysis

Urinary OXT samples were collected before drug administration and after the interaction session, immediately stored at −80°C, and shipped to the University of Hawaii Cancer Center (Honolulu, HI, USA) for biochemical analysis. Additional extraction and assay details are provided in S1 Appendix, **Supplementary Methods**. Samples with absorbance values outside the standard curve range, or with incomplete pre-post pairs, were excluded from the pharmacological manipulation-check analysis, yielding a final urinary OXT sample of OXT *n* = 29 and PLA *n* = 27.

Urinary OXT concentrations were used as a group-level pharmacological manipulation check. Missing or invalid urinary data did not affect inclusion in behavioral analyses. Samples were not normalized to creatinine or specific gravity, so concentrations were not interpreted as estimates of individual systemic exposure (4.4). Analyses therefore tested whether the groups showed differential pre-to-post change consistent with pharmacological exposure. Exploratory individual-level associations are additionally reported in 3.1; they are uncorrected and not interpreted as evidence about systemic or central exposure.

#### Behavioral analysis

Video recordings of the initial and final free-interaction periods (5 min each) were coded in ThemeCoder (PatternVision Ltd.) by trained coders working to a common set of behavioral definitions agreed in joint training (blinding, 2.3). Predefined affiliative behaviors— looking, touching, smiling, speaking, name vocalization, relocating, toy presentation and praising—were drawn from prior human–dog and human–aibo literature [9–12,43–45], and onsets and offsets were annotated to derive duration and frequency measures. Relocating and toy presentation were coded but observed too infrequently to analyze and are not included in the released data, leaving ten outcomes: the duration and frequency of touch, looking, smiling and speaking, together with praise frequency and name-vocalization frequency. For touch, the mean duration of individual touch episodes was additionally derived—total touch duration divided by the number of coded episodes. During the structured task-learning phase, coding focused on looking and touching; aibo’s task behavior was coded for task initiation and duration. The temporal organization of behavior was characterized with T-pattern analysis [26,27] (S1 Appendix, **Supplementary Methods**).

Each segment was coded by one coder and checked by a second, with disagreements resolved through joint video review, so review coverage was complete rather than sampled; the values analyzed are those consensus codes. K.C.M. and S.A. conducted and supervised the coding. Independent pre-consensus records were not retained, so formal inter-rater reliability cannot be computed and none is reported. The 2.5-min time bins were, however, coded in a pass separate from the 5-min period totals, which permits an internal consistency check on the duration measures: agreement between the summed bins and the independently coded period total was ICC(A,1) = .97 for touch duration in the initial free interaction, the measure carrying the primary finding, and ranged from .87 to .98 for the other duration measures, with the exception of speak duration in the final free interaction (.48). The coding procedure, data screening, the full consistency table, and what can and cannot be inferred in the absence of reliability estimates are documented in S1 Appendix, **Supplementary Methods and Supplementary Results.**

### 2.5. Statistical analysis

All analyses were conducted in Python 3.10 (numpy, scipy, statsmodels, pandas, pyreadstat, scikit-learn; S1 Appendix, Supplementary Methods). Statistical significance was evaluated using two-tailed tests with α = .05. Repeated-measures (within-session) effects were analyzed using linear mixed-effects models (LMMs) with a by-participant random intercept, estimated by maximum likelihood; fixed effects and their interactions were evaluated using likelihood-ratio tests, reported as χ². LMMs were adopted as the primary approach because they use all available observations without listwise deletion—an advantage for the fine-grained temporal analyses, in which some segment-level data were missing. For the complete, balanced two-period comparisons the exact repeated-measures F is reported as the primary test, because the maximum-likelihood likelihood-ratio χ² is anticonservative at this sample size [46]; the mixed-model values are reported alongside it in **3.3,** and partial η² is taken from the corresponding ANOVA (S1 Appendix, **Supplementary Methods**). Between-group comparisons at the initial free interaction used independent-samples *t*-tests, with Cohen’s d as the standardized effect size. Welch-corrected tests are reported alongside the pooled-variance tests for the two outcomes where Levene’s test indicated unequal variances (Levene’s test on mean-centered deviations; smile frequency, *p* < .001; look duration, *p* = .042); neither conclusion changes.

#### Hypothesis-driven analysis 1

Early affiliative engagement and its specificity. We examined all coded affiliative behaviors during the initial free-interaction period—touch, looking, smiling, and speaking (each as duration and frequency), praise frequency, and frequency of speaking the robot’s name—using independent-samples *t*-tests comparing the OXT and PLA groups, with Cohen’s d as the standardized effect size. Rather than designating primary outcomes, we tested the directional prediction that the OXT effect would be concentrated in tactile engagement (touch) and graded across channels, with little or no effect on instrumental behavior (speaking). The temporal organization of behavior was examined separately via T-pattern analysis (2.4).

#### Hypothesis-driven analysis 2

Temporal attenuation of OXT effects. To test whether OXT-associated affiliative engagement attenuated across the session, we fit LMMs to behaviors coded during the initial and final free-interaction periods, each with Treatment (OXT vs. PLA) as a between-subjects fixed effect, Period (initial vs. final) as a within-subjects fixed effect, their interaction, and a by-participant random intercept. The confirmatory family comprised the four duration measures (touch, look, smile, speak), to which Holm–Bonferroni correction was applied; frequency, praise, and name-vocalization outcomes were treated as secondary. The Period × Treatment interaction was the effect of interest; significant interactions were decomposed with protected simple-effects tests—independent-samples *t*-tests comparing groups at each period and paired-samples *t*-tests examining within-group change across periods. To characterize the within-session trajectory, each 5-min period was split into two 2.5-min bins (T1–T2 initial; T3–T4 final) and modeled with Time and Treatment (S1 Appendix, Supplementary Methods). These analyses were exploratory; interactions with p between .05 and .10 are reported as trends.

#### Exploratory analyses (candidate mechanisms underlying attenuation)

All analyses in this section were exploratory and were conducted to examine candidate processes that might account for attenuation of the initial OXT-associated affiliative effect. Full details and results for exploratory analyses are reported in S1 Appendix, **Supplementary Results**.

#### Multiple comparisons

Because no behavior was designated a priori as a primary outcome, the family of between-group comparisons at the initial free interaction comprised all ten coded affiliative behaviors. Family-wise error was controlled using Westfall–Young step-down maxT permutation correction (20,000 permutations; S1 Appendix, Supplementary Methods). Uncorrected, Holm-adjusted, and Westfall–Young-adjusted *p*-values are reported together (3.2). For the repeated-measures analyses (H2), Holm correction was applied across the four duration outcomes. The narrower family is used there because H2 is a prediction about the magnitude of sustained engagement, which the duration measures operationalize, and the frequency measures do not: a prediction that engagement would be maintained or amplified is a prediction about time spent engaged. Because this differs from the ten-outcome family used for H1, the Holm-adjusted values across all ten repeated-measures outcomes are also reported as a sensitivity analysis (3.3). Exploratory analyses are labeled as such. Effect sizes are reported with 95% confidence intervals.

#### Baseline characteristics

Baseline demographic, personality, empathy, affective, robot-appraisal, and dog-related variables are summarized descriptively using means, standard deviations, and standardized mean differences (SMDs) between groups (**Table 1; S1 Appendix, Table A;** S1 Appendix, Supplementary Methods). Variables with |SMD| > 0.25 were carried forward into covariate-adjusted sensitivity analyses of the primary outcomes: eleven met that threshold, and each was entered individually, followed by a model containing all eleven simultaneously (3.5). The same threshold was applied to the categorical composition of the sample: because the largest single-category imbalance exceeded it, race was entered as an additional adjustment (3.5). Perceived treatment assignment, recorded during the post-interaction phase, is reported as a blinding check (3.5).

## 3. Results

### 3.1. Pharmacological manipulation check

#### Group-level exposure

An LMM on urinary OXT concentrations (pg/mL), with Session (Pre, Post) and Treatment (OXT, PLA) as fixed effects and a by-participant random intercept, revealed no significant main effect of Treatment, χ²(1) = 2.69, *p =* .101, or Session, χ²(1) = 1.16, *p =* .281 **(Fig 2).** However, a significant Session × Treatment interaction was observed, χ²(1) = 18.37, *p* < .001, indicating differential pre-to-post changes across treatment conditions.

**Fig 2.**
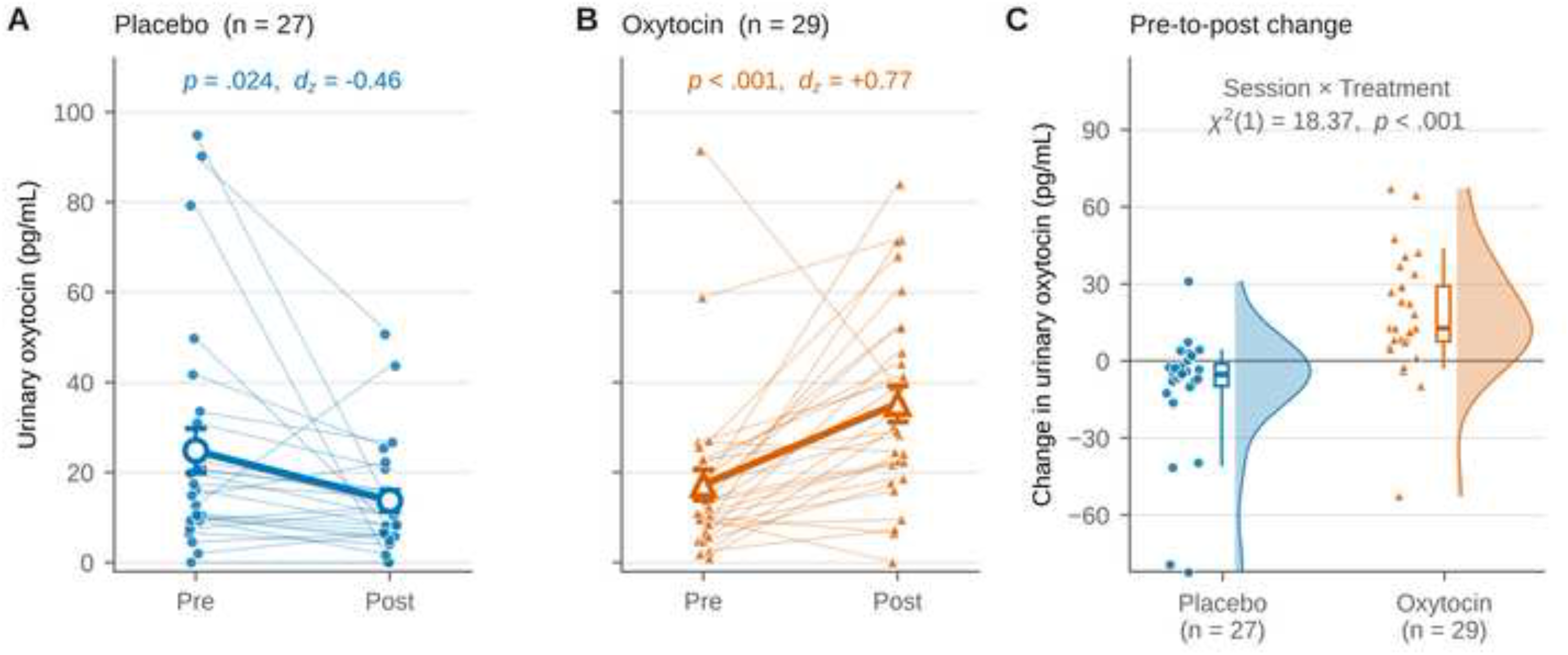
Urinary oxytocin concentrations confirmed group-level pharmacological exposure. (A, B) Individual pre- to post-interaction concentrations. Urinary OXT concentrations (pg/mL) for participants with complete paired samples in the PLA (A; *n* = 27) and OXT (B; *n* = 29) groups. Thin lines connect repeated measurements within participants; the heavy line shows the group mean with standard errors. Concentrations increased in the OXT group, *t*(28) = 4.14, *p* < .001, *d*_z_ = +0.77, and decreased in the PLA group, *t*(26) = −2.39, *p* = .024, *d*_z_ = −0.46; the PLA decrease is not interpreted as a treatment-related effect and may reflect diurnal variation, hydration-related dilution, or other non-specific sampling factors. (C) Pre-to-post change scores by group. Clouds show kernel densities, points show individual participants, and boxes show the median and interquartile range with whiskers at the 10th and 90th percentiles. A linear mixed-effects model yielded a significant Session × Treatment interaction, χ²(1) = 18.37, *p* < .001, with no main effect of Treatment, χ²(1) = 2.69, *p* = .101, or Session, χ²(1) = 1.16, *p* = .281. Groups did not differ at baseline, *t*(57) = −1.37, *p* = .175 (OXT *n* = 29, PLA *n* = 30), and differed at the post-interaction sample, Welch *t*(45.3) = 4.50, *p* < .001. Sample sizes differ across analyses because of incomplete sampling (baseline *n* = 59, post-interaction *n* = 58, complete pairs *n* = 56; S1 Appendix, Table B). Urine was collected approximately 60 min after administration and was not normalized to creatinine or specific gravity; concentrations are interpreted only as a group-level manipulation check. Distributions are right-skewed and are shown on a linear scale because several values are zero. OXT, oxytocin; PLA, placebo.

Groups did not differ in baseline OXT concentrations, *t*(57) = −1.37, *p =* .175, *d* = −0.36. Post-interaction concentrations were higher in the OXT group, *t(45.3)* = 4.50, *p* < .001. Within-group comparisons showed a pre-to-post increase in the OXT group, *t*(28) = 4.14, *p* < .001, *d*_z_ = +0.77, and a decrease in the PLA group, *t*(26) = −2.39, *p =* .024, *d*_z_ = −0.46 (Fig **2**), the latter not interpretable from these data: without normalization to creatinine or specific gravity, a decrease of this size cannot be separated from diurnal variation, hydration, or sampling factors (4.4). Urinary OXT is therefore interpreted only as a group-level manipulation check.

#### Individual-level associations

Exploratory analyses examined whether urinary OXT was associated with tactile behavior at the individual level. No association survived correction for the fifteen tests reported, and these analyses therefore provide no dependable evidence of an individual-level concentration–response relationship, consistent with the use of urinary OXT solely as a group-level manipulation check (**S1 Appendix, Table D; S1 Appendix, Fig B**).

### 3.2. Hypothesis-driven analysis 1: Early affiliative engagement

#### Multivariate effect of treatment

First, it was tested whether treatment acted on the duration outcomes as a family. Four omnibus procedures were applied, and they do not agree, in a way that is itself informative. Of the three that aggregate across all four outcomes, the directional sum of signed effect sizes was significant, *p =* .023, while the parametric MANOVA and its assumption-free analogue fell just short, Wilks’ Λ = .854, *F*(4, 56) = 2.40, *p =* .061, and a permuted Hotelling’s T², *p =* .056; the fourth, the permuted maximum absolute effect size, was significant at *p =* .009, but it re-expresses the family-wise-corrected test for the largest outcome rather than adding independent evidence (S1 Appendix, Supplementary Methods). Across the six frequency outcomes no test reached significance, Λ = .867, *F*(6, 54) = 1.38, *p* = .240, all permutation tests *p* ≥ .053. The pattern—significant when effects are summed directionally, null when power is spread evenly across four outcomes—is what an effect concentrated in one channel and consistent in sign across the rest should produce, and it anticipates the per-outcome results below (**S1 Appendix, Table E**).

#### Sustained tactile engagement

Within the duration family, the effect was concentrated in tactile contact. During the initial free interaction, OXT participants spent more time touching aibo (*M* = 111.53 s, *SD* = 50.42, *n* = 30) than PLA participants (*M* = 77.26 s, *SD* = 37.55, *n* = 31), *t*(59) = 3.02, *p =* .004, *d* = 0.77, 95% CI [0.25, 1.29]. This was the largest effect among all coded behaviors and the only one to survive family-wise correction across the ten-behavior family (Westfall– Young-adjusted *p =* .024; Holm-adjusted *p =* .038).

The difference reflected a shift in the whole distribution rather than a few high-touch participants. The groups separated at every decile from the 10th (52.6 s vs. 31.0 s) to the 90th (157.5 s vs. 116.0 s), and 80% of OXT participants exceeded the PLA median (**Fig 3A; S1 Appendix, Table F**).

**Fig 3.**
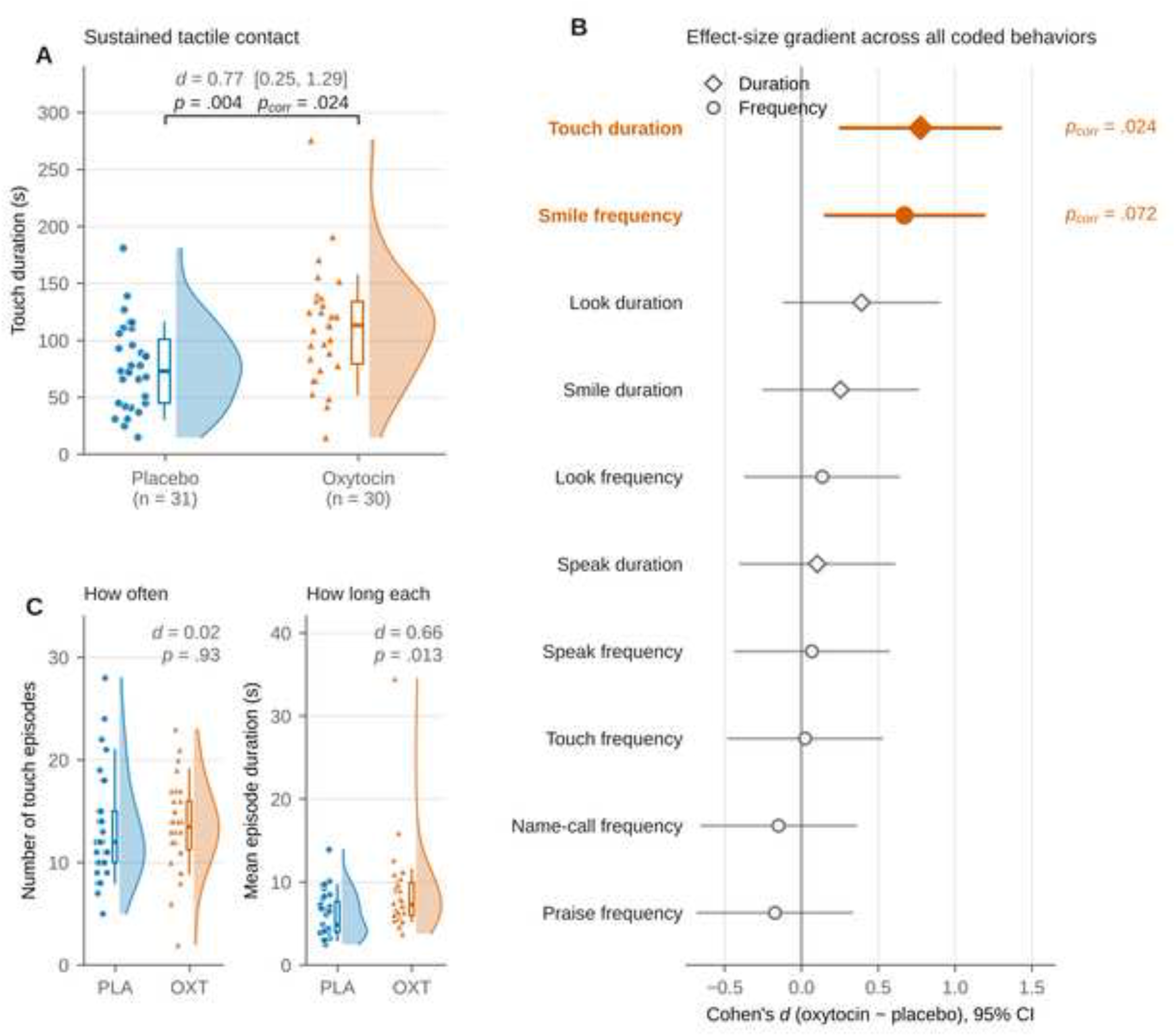
Oxytocin prolonged sustained tactile engagement with aibo during the initial free interaction. (A) Touch duration by treatment group. Clouds show the kernel density of the observed distribution, points show individual participants, and boxes show the median and interquartile range with whiskers at the 10th and 90th percentiles. The OXT group touched aibo longer than the PLA group, *t*(59) = 3.02, *p* = .004, *d* = 0.77, 95% CI [0.25, 1.29]; *p* = .024 after Westfall–Young step-down correction across all ten coded behaviors. The difference was a distributional shift rather than an outlier effect: the groups separated at every decile from the 10th to the 90th percentile, and the result held under non-parametric, permutation, trimmed, winsorized, and log-transformed tests (S1 Appendix, Table F). (B) Effect-size gradient across all coded behaviors. Standardized mean differences (Cohen’s d, OXT − PLA) with 95% confidence intervals for all ten coded affiliative behaviors, ordered by effect size; diamonds denote duration outcomes and circles denote frequency outcomes. Touch duration was the only behavior to survive family-wise correction; smile frequency was trend-level (*p* = .072). Across the four duration outcomes the directional sum of signed effect sizes was significant, *p* = .023, while the parametric MANOVA and the permuted Hotelling’s T² fell just short, Wilks’ Λ = .854, *F*(4, 56) = 2.40, *p* = .061, and *p* = .056; no omnibus test was significant across the six frequency outcomes, *F*(6, 54) = 1.38, *p* = .240 (S1 Appendix, Table E). (C) Decomposition of the touch-duration effect. Groups did not differ in the number of touch episodes initiated (*d* = 0.02, *p* = .93; equivalence supported within ±0.5 *SD*, TOST *p* = .034), but episodes lasted longer under OXT (8.84 s vs. 5.98 s; *d* = 0.66, *p* = .013). The 1.44-fold group difference in total touch duration was therefore accounted for by a 1.48-fold difference in mean episode duration rather than by more frequent contact. *n* = 30 (OXT) and 31 (PLA) throughout. CI, confidence interval; MANOVA, multivariate analysis of variance; OXT, oxytocin; PLA, placebo; pcorr, Westfall–Young-corrected p; TOST, two one-sided tests.

#### Specificity across behavioral channels

The remaining behaviors formed a graded falloff. A convergent but weaker effect appeared for positive affective display: the OXT group smiled more frequently (*M* = 4.77, *SD* = 4.65) than the PLA group (*M* = 2.29, *SD* = 2.44), *t*(59) = 2.62, *p* = .011, *d* = 0.67, 95% CI [0.15, 1.19]; Welch-corrected, *t*(43.5) = 2.59, *p* = .013. This did not survive family-wise correction (corrected *p* = .072) and is interpreted as suggestive. Looking and smiling duration showed small, non-significant differences (*d* = 0.39 and 0.25; look duration Welch-corrected, *t*(53.0) = 1.53, *p* = .131), whereas speaking, praise, name vocalization, and touch frequency showed essentially no group difference (all |*d*| ≤ 0.18; **Fig 3B; S1 Appendix, Table G**). OXT did not alter the temporal organization of behavior (**S1 Appendix, Table H**).

#### Sustained contact, not initiation

The two groups did not differ in how often touch was initiated: OXT participants made no more touch episodes (*M* = 13.50, *SD* = 4.42) than PLA participants (*M* = 13.39, *SD* = 5.26), *t*(59) = 0.09, *p* = .928, *d* = 0.02, 95% CI [−0.48, 0.53]; two one-sided tests supported equivalence within a margin of ±0.5 *SD*, *p* = .034, so this is an informative null rather than an absence of evidence; the margin is a conventional small effect and was not specified in advance. Episodes instead lasted longer under OXT (*M* = 8.84 s, *SD* = 5.55) than under PLA (*M* = 5.98 s, *SD* = 2.74), *t*(59) = 2.57, *p* = .013, *d* = 0.66, 95% CI [0.14, 1.17]; the same held non-parametrically (*p* = .004) and after log transformation (*p* = .001; S1 Appendix, Supplementary Results). OXT therefore prolonged episodes of contact already under way rather than prompting participants to reach for the robot more often (**Fig 3C**). This decomposition was derived during analysis and is exploratory; the episode measure is not a member of the corrected ten-outcome family, and its *p*-value is uncorrected. The early effect was thus concentrated in sustained tactile engagement, graded across channels, absent for instrumental behavior, and specific to the magnitude rather than the structure of behavior—the pattern H1 anticipated, although only the first of these four respects rests on a corrected significant test, the remaining three being null-based and therefore compatible with, rather than confirmatory of, the prediction.

### 3.3. Hypothesis-driven analysis 2: Temporal attenuation of OXT effects

#### Period × Treatment effects

Contrary to the consolidation prediction (H2), the early OXT advantage did not persist. Repeated-measures models comparing the initial and final free-interaction periods revealed a significant Period × Treatment interaction for touch duration, *F*(1, 59) = 6.79, *p* = .012, η²ₚ = .103, and for looking duration, *F*(1, 59) = 5.86, *p* = .019, η²ₚ = .090, with smiling duration at trend level, *F*(1, 59) = 3.30, *p* = .074. After Holm correction across the four duration outcomes, only touch duration remained significant (*p* = .046); looking duration did not (*p* = .056), nor did smiling (*p* = .149). The looking result also differs from touch in kind: the groups did not differ at either period (initial *d* = 0.39, *p* = .133; final *d* = −0.29, *p* = .256), so the interaction reflects a crossover between two differences that were never established rather than the attenuation of a demonstrated advantage, and we do not interpret it further. The mixed-model likelihood-ratio values, under which looking duration reaches *p* = .049 after correction, are in S1 Appendix, Table I; that difference is why the exact *F* is primary here (**2.5**). Because H1 used a ten-outcome family, we also report the corresponding sensitivity analysis: correcting across all ten repeated-measures outcomes, touch duration does not survive (*p* = .116). The attenuation result therefore depends on treating the four duration measures as the confirmatory family for H2, as specified in 2.5, and should be read with that dependence in view (**S1 Appendix, Table I**).

#### Simple effects

Simple-effects tests showed that the groups converged through movement in both directions. Touch duration declined in the OXT group (111.53 s to 91.87 s), *t*(29) = −2.20, *p =* .036, *d*_z_ = −0.40, and increased non-significantly in the PLA group (77.26 s to 90.32 s), *t*(30) = 1.48, *p =* .149, *d*_z_ = +0.27. For looking and smiling the OXT group declined (*d*_z_ = −0.69 and −0.44) while the PLA group changed less (*d*_z_ = −0.06 and −0.30; S1 Appendix, Table I). Convergence in tactile engagement therefore reflected a decline in the OXT group together with a smaller, non-significant increase in the PLA group. By the final interaction no group difference remained for touch, looking, or smiling (**all *p* ≥ .256; Fig 4A–C**).

**Fig 4.**
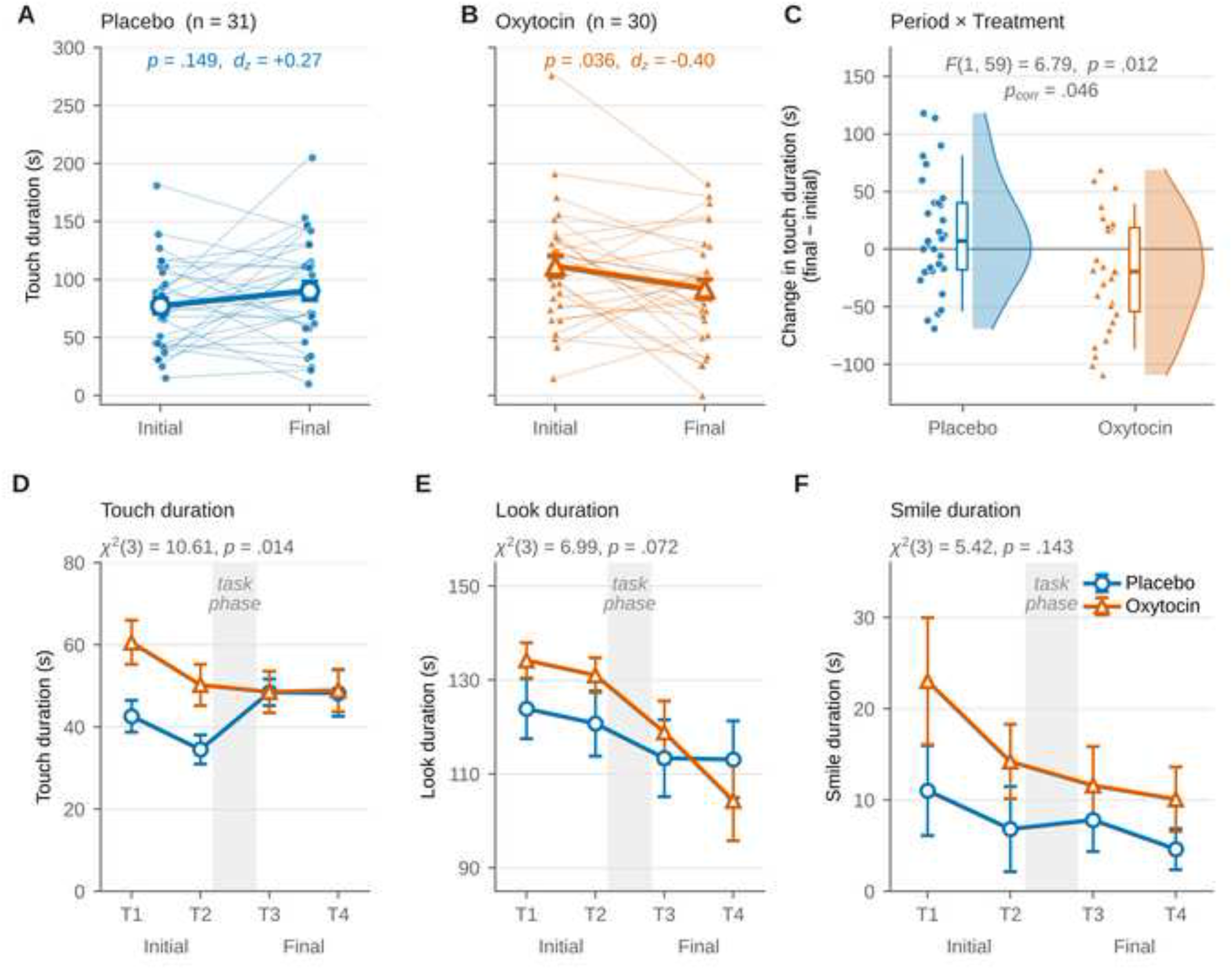
The early oxytocin effect attenuated within the session. (A, B) Touch duration across the two free interactions. Individual trajectories for the initial and final free interactions in the PLA (A) and OXT (B) groups. Thin lines connect repeated measurements within participants; the heavy line shows the group mean with standard errors. Touch duration declined in the OXT group, *t*(29) = −2.20, *p* = .036, *d*_z_ = −0.40, and increased non-significantly in the PLA group, *t*(30) = 1.48, *p* = .149, *d*_z_ = +0.27; convergence by the final interaction therefore reflected movement in both groups rather than decline in the OXT group alone. Groups did not differ at the final interaction, *t*(59) = 0.13, *p* = .895. **(C) Period × Treatment change scores.** Change in touch duration (final − initial) by group. The Period × Treatment interaction was significant, *F*(1, 59) = 6.79, *p* = .012, *p* = .046 after Holm–Bonferroni correction across the four duration outcomes. **(D–F) Fine-grained trajectories across the four 2.5-min bins.** Group means with standard errors for touch (D), look (E), and smile (F) duration. T1 and T2 divide the initial free interaction, T3 and T4 divide the final free interaction, and the shaded band marks the intervening structured task phase. The Time × Treatment interaction was significant for touch duration, χ²(3) = 10.61, *p* = .014, a non-significant trend for look duration, χ²(3) = 6.99, *p* = .072, and non-significant for smile duration, χ²(3) = 5.42, *p* = .143. Both groups declined across the initial interaction by comparable amounts (touch duration, OXT *d*_z_ = 0.43, PLA *d*_z_ = 0.50); what distinguishes them is that the group difference closed across the intervening task phase. These analyses were exploratory. Interaction period is necessarily confounded with elapsed time, loss of novelty, task-related reframing, and the pharmacokinetic time course of intranasal OXT; the pattern is therefore described as within-session attenuation. OXT, oxytocin; PLA, placebo; pcorr, Holm–Bonferroni-corrected p.

Mean touch-episode duration showed a Period × Treatment interaction at the threshold of significance, *F*(1, 59) = 3.76, *p =* .057: episodes lengthened in the PLA group (5.98 s to 7.70 s, *p =* .038) while OXT episodes were unchanged (8.84 s to 8.16 s, *p =* .483), so at this level convergence was produced mainly by the PLA group catching up.

These findings did not support the consolidation prediction; they revealed attenuation of the early OXT-associated advantage across the session (**Fig 4; S1 Appendix, Table I**).

#### Fine-grained trajectory

Fine-grained LMMs, fit to all available segment-level observations, examined the trajectory across the four temporal bins (T1–T4). For touch duration, the Time × Treatment interaction was significant, χ²(3) = 10.61, *p =* .014 (S1 Appendix, **Supplementary Results**).

Thus, the fine-grained analyses provided exploratory support for a differential OXT-related trajectory in touch duration **(Fig 4D**) and weaker, trend-level evidence for looking duration (**Fig 4E**), but no evidence for a differential trajectory in smile duration (**Fig 4F**). The bin-level comparisons locate when the difference closed. Touch duration declined in both groups across the initial interaction, in parallel: the group difference was 18.0 s in the first 2.5-min bin, *t*(59) = 2.73, *p* = .008, *d* = 0.70, and 15.7 s in the second, *t*(59) = 2.58, *p* = .012, *d* = 0.66. In the final interaction it was 0.1 s and 0.7 s (both *p* ≥ .93). The early advantage therefore held throughout the initial interaction and closed across the intervening task phase (S1 Appendix, **Supplementary Results**). Because interaction period is necessarily confounded with elapsed time, loss of novelty, task-related reframing, and the pharmacokinetic time course of intranasal OXT, this pattern is described as within-session attenuation and is not interpreted as evidence about consolidation as a psychological process.

### 3.4. Exploratory analyses of candidate mechanisms underlying attenuation

All analyses in this section were exploratory and are reported in full in S1 Appendix, Supplementary Results. During the cooperative task, OXT participants touched aibo more than PLA participants, χ²(1) = 4.60, *p* = .032 (exploratory, uncorrected), a difference present in the first two trials and absent in the third, which requires touching aibo to complete (**S1 Appendix, Fig C, panel A)**, and trick-learning success neither predicted nor moderated the affiliative decline (**Table 2**). Three further candidates did not account for the attenuation: OXT did not alter the structural organization of behavior, change in robot appraisal did not predict behavioral change, and the primary effects survived adjustment for baseline appraisal (S1 Appendix, Tables H, J and K). Robot appraisal was not wholly unaffected by treatment, however: Positive Social Attribution rose from pre- to post-interaction, χ²(1) = 6.88, *p =* .009, and did so differentially by group, χ²(1) = 4.48, *p =* .034. This effect is exploratory and uncorrected, it does not mediate the behavioral findings, and we report it because a treatment effect on a self-reported outcome should not be left to the supplement.

**Table 2.**
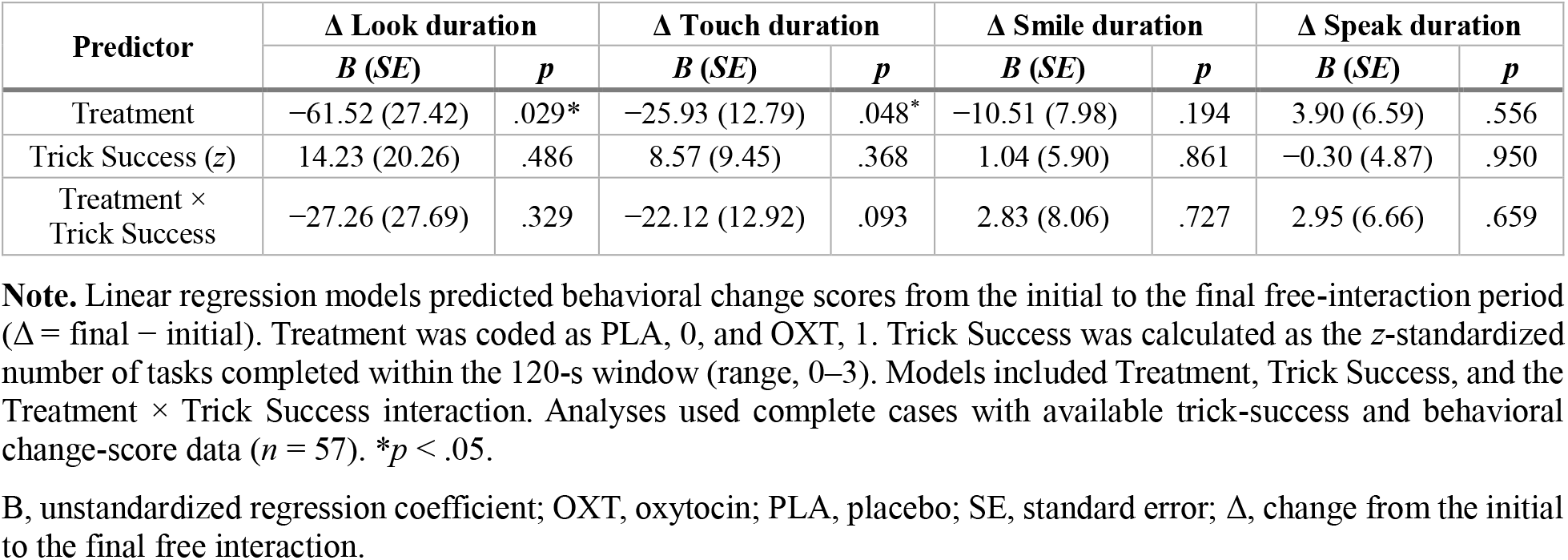
Regression models predicting change scores in affiliative behavior from treatment and trick-learning success.

### 3.5. Baseline characteristics and blinding

#### Baseline imbalances

Baseline characteristics are reported descriptively in **Table 1** and **S1 Appendix, Table A**. Eleven variables showed standardized mean differences exceeding |0.25|. Each of the eleven was entered individually as a covariate, and all eleven were then entered simultaneously. The treatment effect on touch duration remained significant in every single-covariate model (*B* = 26.14 to 38.80, all *p* ≤ .023) and with all eleven entered, *B* = 39.05, *p* = .041, as did mean touch-episode duration (*p* ≤ .042 individually; jointly *B* = 2.92, *p* = .024). Smile frequency survived each single-covariate model (*p* ≤ .026) but not simultaneous adjustment (*B* = 2.26, *p* = .195), consistent with its status as a suggestive rather than an established effect (**S1 Appendix, Table K**). The sample also differed in racial composition (**Table 1**), which is relevant because aibo is a culturally salient product and attitudes toward robot dogs vary cross-culturally [37]. Adjusting for race left the treatment effects unchanged: touch duration, B = 35.73, p = .003; smile frequency, B = 2.56, p = .012; mean touch-episode duration, B = 2.88, p = .015 (S1 Appendix, Supplementary Results).

#### Blinding check

Blinding was assessed by asking participants at the end of the session which treatment they believed they had received. Perceived assignment did not differ between groups: 36.7% of OXT participants and 25.8% of PLA participants believed they had received OXT, χ²(1) = 0.84, *p =* .360, indicating that participants were not able to identify their treatment condition above chance.

### 3.6. Control measures

In LMMs, heart rate and negative affect decreased over time, consistent with laboratory habituation, but no significant Treatment or Time × Treatment effects emerged for any physiological or mood measure, suggesting that the behavioral findings were not attributable to generalized changes in arousal or affective state (**S1 Appendix, Table L**). No adverse events, withdrawals, or dropouts occurred.

## 4. Discussion

The present study tested whether intranasal OXT enhances affiliative engagement toward a socially ambiguous artificial agent, and whether such engagement persists across an evolving interaction. H1 was supported in its central respect: the early effect was present and was concentrated in tactile engagement. H2 was not: the advantage was not consolidated across the session. Three findings stand out. First, OXT produced a behaviorally specific early effect: the omnibus tests across the duration family were mixed and did not establish a distributed multivariate effect, and the effect was instead concentrated in sustained tactile contact, graded across channels, and absent for instrumental behavior. Second, the effect was on the maintenance of contact rather than its initiation: OXT participants did not reach for the robot more often, but each episode of contact lasted longer. Third, the early advantage did not persist. The groups converged by the final interaction, and the fine-grained analyses locate that convergence across the intervening task phase. Together these results suggest that OXT transiently amplified the salience of an early, novel social encounter rather than initiating an affiliative bond that consolidates with continued interaction, supporting context-sensitive, salience-based accounts of OXT function rather than a uniformly prosocial or bond-building role [1–3].

### 4.1. Early affiliative engagement: Sustained tactile contact under social ambiguity

#### Shape of the effect

The early OXT effect was specific, and its shape is informative. Participants under OXT initiated contact no more often than participants under PLA—the number of touch episodes was essentially identical—but each episode lasted longer, and the difference in total contact time was almost entirely accounted for by that difference. OXT did not make participants reach for the robot more readily; it made them keep their hands on it once they had. This decomposition was derived during analysis rather than predicted and is offered as the most informative description of the effect rather than as a confirmed hypothesis.

#### Affective touch

This dissociation aligns the finding with work on affective touch. Intranasal OXT increases the perceived pleasantness of social touch and accompanying neural responses [47], increases the pleasantness of affective touch together with orbitofrontal cortex activity [48], and does so selectively for gentle, C-tactile-targeted stroking rather than moderate-pressure contact [49]. If OXT raises the moment-to-moment hedonic value of tactile contact, the behavior it should change is how long contact is maintained rather than how often it begins, which is what we observe.

#### Qualifications

Two qualifications limit this interpretation. The affective-touch studies concern touch received from another person, whereas our participants were the agents of contact toward a partner that could not reciprocate in kind; whether the same mechanism governs sustained self-initiated touching is an open question rather than an established extension (S1 Appendix, **Supplementary Discussion**). And we did not measure the perceived pleasantness of touching aibo, so increased tactile pleasantness remains one candidate among several: heightened tactile salience, curiosity, sensory exploration and novelty-driven engagement all predict longer contact without implying it was more pleasant. Distinguishing them requires measures this design did not collect.

Two features of the effect further constrain its interpretation. OXT altered the magnitude of engagement but not its structural organization, indicating a selective rather than global reorganization of behavior (3.2, 3.4). It was also specific in channel: concentrated in sustained tactile engagement, graded across the remaining behaviors, and absent for instrumental behavior (3.2).

### 4.2. Within-session attenuation: What the design can and cannot separate

#### What the design can separate

The early OXT advantage did not persist; by the final free interaction the groups no longer differed in touch, looking, or smiling duration. Before interpreting this, it matters what the design can and cannot separate. Interaction period was fixed rather than counterbalanced, so the contrast between the initial and final free interactions confounds elapsed time since administration, loss of novelty, the intervening reframing of aibo as a task object, and the pharmacokinetic time course of intranasal OXT itself. The window in which the group difference was detectable began roughly 30 min after administration and lasted five minutes; the final free interaction fell roughly an hour after administration. Given continuing uncertainty about the time course of behaviorally relevant central OXT exposure after intranasal delivery [19,21–24], we cannot exclude that some or all of the attenuation reflects declining pharmacological exposure rather than a psychological process. We therefore describe the finding as within-session attenuation and do not treat it as evidence about consolidation as a psychological mechanism.

#### How the attenuation arose

Two observations bear on how the attenuation arose. The first is the timing. The group difference held through the whole of the initial free interaction—both groups declined across it in parallel, and the difference was still significant in its second half—and had closed by the first half of the final interaction. Convergence therefore coincided with the intervening task phase, which leaves the reframing of aibo as a task object a live candidate rather than one these data exclude. Trick-learning success neither predicted nor moderated the decline, which argues against the strongest form of that account, in which success at the task is what displaced affiliative contact; it does not rule out the weaker form, in which the task changes what the robot is for. The second is that convergence was not simply the OXT group falling back: for total touch duration the OXT decline and the non-significant PLA increase were of comparable magnitude, while at the level of episode duration the OXT group was unchanged and the PLA group lengthened. A description in which OXT front-loads sustained contact that the PLA group arrives at more slowly fits these data better than one in which the OXT effect simply decays, though the design cannot decide between them.

#### How H2 failed

A precise statement of how H2 failed is worth making, because the failure is partial. The prediction had two components: convergence by the final interaction, and convergence at a sustained level achieved by the PLA group catching up to a maintained OXT group. The first was observed; the second was not, because total tactile engagement fell in the OXT group. The one measure that did behave as H2 predicted was mean touch-episode duration, which remained flat under OXT while lengthening under PLA—though that interaction was only at the threshold (*p =* .057) and falls outside both confirmatory families. The prediction therefore failed for total contact time but not for the per-episode measure that carried the original effect. Because interaction period is confounded with elapsed time and pharmacokinetics, we cannot attribute this failure confidently to a psychological process such as non-consolidation; what the data establish is that the early advantage did not persist in the form predicted.

#### Regression to the mean

Convergence from different starting points invites a regression-to-the-mean explanation. These groups were formed by randomization, so measurement error alone cannot generate a Period × Treatment interaction; but H2 was examined because H1 was significant, and conditional on a large initial difference some regression toward the mean is expected. We therefore treat the magnitude of the attenuation as less well estimated than its direction, and the analysis-of-covariance form, with the reasons it is not the appropriate estimator here, is reported in S1 Appendix, Supplementary Results.

#### Accounts consistent with the pattern

The pattern is consistent with allostatic and social-salience accounts, on which OXT effects are strongest under novelty and uncertainty and diminish as a context becomes predictable [3,50]. It is also consistent with a feature of the partner: in dog–human interaction reciprocal gaze supports OXT-linked positive feedback [11,51], and dogs elicit stronger companionship responses than aibo [12]. A robot that responds contingently but does not reciprocate affiliation in kind may support the initiation of affiliative behavior without supplying the signals that sustain it (4.4). Whether attenuation reflects limited reciprocity rather than habituation to novelty therefore remains open, and the two are not mutually exclusive.

### 4.3. Exploratory analyses and unresolved mechanisms

The exploratory analyses constrain but do not identify a mechanism: behavioral adaptation, trick-learning success, behavioral-sequence complexity, and explicit robot appraisal did not account for the attenuation. The timing is at least consistent with implicit contextual updating—the difference closed across the task phase, and participants may have shifted from treating aibo as a potential social partner to treating it as a task object—but the available measures cannot test that account, and elapsed time and pharmacokinetics are confounded with it.

These exploratory analyses were modestly powered, and the measures were indirect; the null results constrain simple interpretations rather than excluding the mechanisms they tested.

### 4.4. Limitations and future directions

Six limitations qualify what these findings can support; we list them in descending order of consequence for the interpretation offered above; how the first three interact, and what would resolve each, are set out in S1 Appendix, Supplementary Discussion.

The first and most consequential limitation is the absence of a comparison condition. Every participant interacted with aibo and only with aibo. Without a human partner, a live dog, or a non-social object as a comparison, the design cannot establish that the effect is specific to robotic or socially ambiguous agents, that sustained contact reflects affiliation rather than general engagement or exploration, or that the observed time course differs from what would be seen with any novel object. The finding should be read as evidence that OXT prolongs tactile engagement with this class of agent, not as evidence about what is special about it.

Second, the reciprocity of the interaction was not measured. Aibo responded contingently to touch, sound, and visual input, and those responses plausibly shaped participants’ behavior—but the robot produced no retrievable log of its own actions, and robot behavior during the free interactions was not coded from video. We therefore have no participant-level index of how responsive each participant’s aibo was and cannot test whether variation in received responsiveness predicted sustained contact or its decline. Because reciprocal contingency is central to most accounts of how affiliative bonds form and are maintained, this is a substantive gap, and it is a specific reason why these data cannot adjudicate between habituation to novelty and insufficient reciprocity.

Third, interaction period was confounded with elapsed time and pharmacokinetics (**4.2**). The fixed session order, including the fixed order of the three tasks, preserved the naturalistic structure of the paradigm, but the temporal findings are best read as descriptive of this sequence.

Fourth, the urinary OXT measure was limited: samples were not normalized to creatinine or specific gravity, a single post-interaction sample was collected at a time point that may fall after peak concentrations, urine was used rather than saliva or plasma, and peripheral concentrations are at best a weak proxy for central activity [19,21,22,52–54]. It is therefore used only to confirm group-level differences in pharmacological exposure, and exploratory within-group analyses showed no relationship with tactile behavior.

Fifth, the study carries the usual constraints of a single-session pharmacological design. Participants were healthy adult males, and OXT effects can differ by sex [1,55]. The design could reliably detect only large between-group effects (**2.4**), so the estimates are imprecise, the interaction and exploratory analyses were underpowered, and the null results are inconclusive. The study was neither preregistered nor entered in a clinical-trial registry, and no primary outcome was designated in advance. Inter-rater reliability could not be computed; blind coding (**2.3**) and the close agreement of the separately coded time bins for touch (**2.4**) limit, but do not remove, this concern.

Finally, the testing environment introduced dependencies the analysis can only partly address. Up to five participants were tested simultaneously in a partitioned room, and five aibo units were in use for participant sessions. Unit identity was recorded and modeling it leaves the findings unchanged (S1 Appendix, **Supplementary Results)**; testing session was not recorded, however, so shared session effects—a common experimenter, time of day, audible neighbors—cannot be estimated. The units are adaptive and were initialized once before the study rather than before each participant, and touch duration did increase across the testing period, in both groups and independently of unit; treatment was balanced across that period and the effect survives adjustment for it, but the source of the trend cannot be identified from these data (S1 Appendix, Supplementary Results).

Despite these constraints, the study characterizes the temporal dynamics of OXT-modulated tactile engagement toward a social robot with a fully reported analysis, individual-level data, and robustness checks, and it identifies a specific behavioral signature—prolonged episodes of contact without more frequent contact—that future work can test directly.

### 4.5. Conclusion

Intranasal OXT transiently prolonged sustained tactile engagement with a socially ambiguous robotic agent during a first encounter. The effect was concentrated in how long contact was maintained rather than how often it was initiated and was no longer detectable by the end of the session. What the study can establish is bounded by its design, as set out above: these findings are not evidence about bond formation, robot-specific mechanisms, or consolidation as a psychological process. They do show that a neuropeptide implicated in mammalian affiliation can measurably alter how people physically engage with an artificial agent within a narrow early window, and they define what the next study will need.

## Acknowledgments

The authors thank all study participants for their time and commitment. The authors also acknowledge the assistance of research staff and student assistants who supported participant scheduling, data collection, and study coordination.

Generative artificial-intelligence tools were used in the preparation of this manuscript, and their role is declared here in full. OpenAI ChatGPT (GPT-5.5) was used for language editing, structural organization, and consistency checking. Anthropic Claude (Opus 5) was used to implement the analysis scripts—including the permutation-based multiple-comparison correction, the omnibus multivariate tests, the robustness battery, the covariate-adjusted sensitivity models, the derived episode-duration measure, and the exploratory urinary-oxytocin correlations—to generate Figs 2–4 and S1 Appendix, Figs A–C, and to draft manuscript text subsequently revised by the authors. All analyses were run against the de-identified dataset, and the analysis and figure-generation scripts are provided with this submission so that every reported value can be independently verified. No AI tool was used to generate, alter, or fabricate research data. All analyses, interpretations, and editorial decisions were reviewed, verified, and approved by the authors, who take full responsibility for the content of this manuscript.

## Supporting information

**S1 Appendix. Supplementary methods, results, tables and figures.**

**S1 Checklist. CONSORT 2025 checklist, used as a reporting aid for this randomized design.**

